# Inference of clonal hematopoiesis using the collective behaviour of DNA methylation states

**DOI:** 10.1101/2025.11.18.689122

**Authors:** Samuel. J. C. Crofts, Caleb M. Grenko, Neil A. Robertson, Kristina Kirschner, Tamir Chandra

## Abstract

Clonal expansion occurs when descendants of a single progenitor cell come to dominate a tissue. We show that this process generates a predictable collective behaviour in DNA methylation landscapes: at CpG sites that faithfully transmit their allelic methylation status across cell divisions, methylation proportions shift toward discrete values (0%, 50%, or 100%) as clones expand. We demonstrate this phenomenon in clonal hematopoiesis, in which a hematopoietic stem cell acquires a fitness-conferring mutation that leads to clonal expansion. Exploiting these dynamics, we developed COMET (Clonal Observation from METhylation), which quantifies clonal burden from bulk methylation data without prior knowledge of driving mutations. Validation against targeted sequencing demonstrated robust prediction across mutation types, accurate longitudinal tracking, and detection of sequencing artifacts. Applied to 15,900 individuals, COMET-predicted clonal hematopoiesis replicated known genetic associations (TCL1A, NRIP1) and phenotypic relationships with smoking and chronic obstructive pulmonary disease. These findings reveal fundamental principles of how clonal dynamics manifest epigenetically and establish a mutation-agnostic approach for clonal burden assessment.

## Main

Hematopoietic stem cells (HSCs) maintain blood production through balanced self-renewal and differentiation. Some somatic mutations shift this equilibrium toward self-renewal, leading to clonal expansion beyond what would be expected from stochastic drift. This phenomenon is generally termed clonal hematopoiesis (CH) and is known as clonal hematopoiesis of indeterminate potential (CHIP) when occurring without overt hematologic pathology^1^.

CHIP predominantly involves mutations in leukemic driver genes^1,2^. It is associated with increased risk of hematologic malignancy^3^, cardiovascular disease^4,5^, poor prognoses in chronic heart failure^6^, and mortality^3^. CHIP’s high prevalence in elderly patients^2,3,7,8^, its increase following genotoxic therapy^9^, and its suspected role in therapy-related secondary leukemias^10^ have made it a research priority.

Gold-standard detection methods rely on genome resequencing; however, distinguishing true somatic drivers from neutral mutations, developmental variants, and sequencing artifacts requires stringent filtering algorithms^11^. We have previously demonstrated how longitudinal sampling, combined with modelling growth trajectories, can reduce the number of false calls and increase sensitivity^12^. Additionally, expert manual curation can further improve accuracy, but cannot eliminate false negatives and false positives^13^.

Due to these challenges, error-corrected gene panel sequencing remains costly and difficult to implement at scale. Methylation-based CH detection offers a cost-effective alternative, though defining general methylation signatures is challenging due to diverse driver mutations creating distinct epigenetic footprints.

Strategies to infer clonality based on the collective behaviour of fluctuating CpGs have recently demonstrated the potential of such approaches^14,15^. Here, we uncovered a phenomenon that emerges from the collective behaviour of a different set of CpGs that are established heterogeneously - but predictably - during development and stably transmit their allelic methylation states throughout life. We show that this phenomenon results in distributions of methylation values driven by the level of clonal expansion in an individual. In addition to presenting the theoretical basis for this phenomenon, we demonstrate its manifestation in human and mouse blood samples, and develop COMET (Clonal Observation from METhylation), an algorithm that exploits these principles to infer CH from bulk DNA methylation data. COMET can track clonal growth longitudinally and detect errors arising from sequencing-based CH detection. Finally, we validate our method by estimating variant allele frequencies (VAFs) in a large external cohort (N = 15,900). We show that estimated VAF is associated with lifestyle factors and health outcomes known to be associated with CH. Additionally, we perform a genome-wide association analysis, which implicates known CH-associated variants.

## Results

### Clonality in the HSC pool can be observed through bulk DNA methylation

When cells undergo clonal expansion, the descendants of a single progenitor come to dominate a tissue. A fundamental question arises: how does this clonal process manifest in the epigenetic landscape? We reasoned that at CpG sites that faithfully transmit their allelic methylation status across cell divisions, clonal expansion should generate a predictable collective behaviour.

We began with the observation that at any given CpG site, a single cell’s methylation status will be one of three allele combinations: both alleles unmethylated (UU), one allele methylated and one unmethylated (MU), or both alleles methylated (MM) (**Fig. 1a**, “Background”). From this basis, we reasoned that if two conditions are met at certain CpG sites, clonal expansion should generate a predictable collective behaviour.

**Figure 1.**
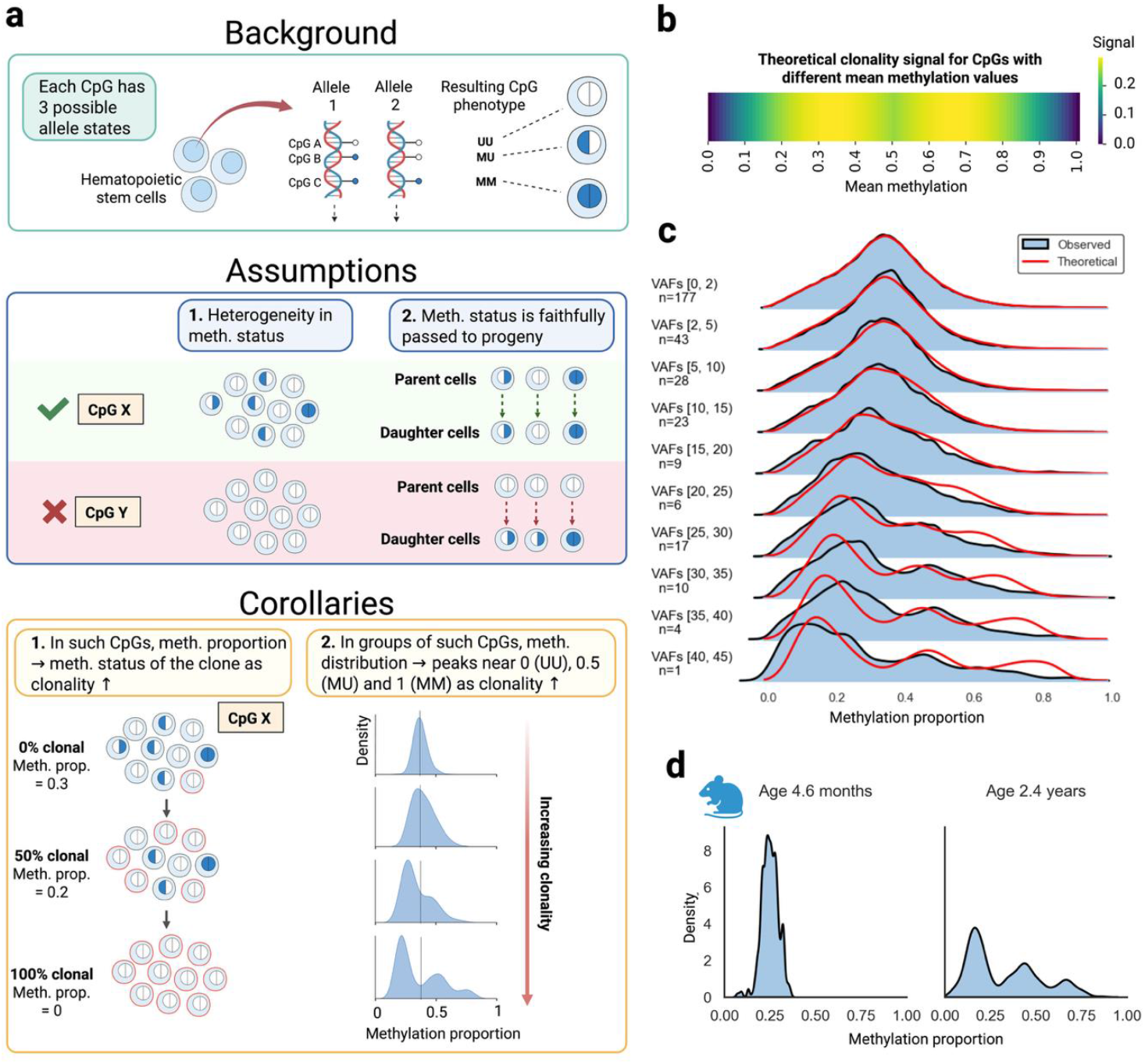
Clonal dynamics can be inferred from the collective behaviour of CpGs. **a**, Overview of the theoretical basis of COMET. Each hematopoietic stem cell (HSC) has two alleles, each of which is either methylated or unmethylated at each CpG, resulting in three allele combinations: UU (unmethylated/unmethylated), MU (methylated/unmethylated, or vice versa), or MM (methylated/methylated). If there is a subset of CpGs which 1) show heterogeneity in allele status within a sample, and 2) faithfully pass this allele status between cellular generations, then: 1) In such CpGs, the measured methylation proportion will increasingly reflect the methylation status of the clone as it expands (i.e. move towards 0, 0.5, or 1, depending on the allele state of the clone) and 2) In groups of such CpGs, peaks will emerge towards 0, 0.5, and 1 as clonality increases and each individual CpG moves closer to 0, 0.5, or 1. Further, the ratio of peak heights will reflect the original underlying allele distribution in this group of CpGs. **b**, Theoretical clonal signal strength across CpGs of different population means if the underlying allele states are distributed according to Hardy-Weinberg equilibrium (see Methods for details). **c**, Observed methylation distributions (blue) of selected CpGs for participants in the Lothian Birth Cohorts binned by error-corrected VAF. Theoretical distributions are shown for each bin (red line, see Methods for details). **d**, Similar clonal dynamics are seen in the methylation distributions of mice. Plots display histograms of mouse-specific clonal CpGs (see Methods for details), with a young mouse shown on the left and an older mouse shown on the right. Created with BioRender.com.

First, methylation values in bulk data — which reflect the proportion of methylated alleles across a population of cells — deviating from 0%, 50%, or 100% at a given CpG site must indicate allelic heterogeneity across cells. Since homogeneous cellular populations would produce only these discrete values (0%, 50%, or 100%), intermediate percentages necessarily reflect mixed methylation patterns across constituent cells. Such sites are necessary to observe the emergence of clonal patterns (**Fig. 1a**, “Assumptions”, panel 1).

Second, if a subset of CpGs faithfully transmits their allelic methylation status across cell divisions, clonal expansion would shift the average methylation of these CpGs as the relative contributions of individual cells within the population change (**Fig. 1a**, “Assumptions”, panel 2).

These assumptions lead to two testable predictions. First, at individual faithful CpGs, the bulk methylation proportion should shift toward the methylation status of the expanding clone as clonality increases (**Fig. 1a**, “Corollaries”, panel 1). In the case of clonal hematopoiesis — in which blood cells are disproportionately descended from a single HSC — one of three scenarios would be expected at each CpG: methylation proportions moving towards 0% (if the original cell had the CpG state UU), 100% (if the original cell was MM), or 50% (if the original cell was MU). The figure illustrates this phenomenon using a CpG for which the stem cell that acquires the CH mutation (circled in red) has the state UU, resulting in the bulk methylation proportion shifting towards 0 as it expands.

Second, when methylation values are observed across many faithful CpGs in an individual, the distribution of methylation proportions should show peaks emerging at 0, 0.5, and 1 as clonality increases (**Fig. 1a**, “Corollaries”, panel 2). Gabbutt et al.^14^ have previously observed collective CpG dynamics in a different context, describing CpGs with methylation values of ∼50% that were “fluctuating” (i.e., stochastically changing their methylation status).

Importantly, because the methylation status of the original mutated HSC is unknown *a priori*, it is impossible to know whether to expect an increase or decrease in the methylation level of any given CpG. This lack of a consistent directional signal at any given CpG would make it challenging to exploit this phenomenon using machine learning methods.

Next, we explored whether the underlying proportions of allele combinations might follow Hardy-Weinberg Equilibrium (HWE). That is, for a CpG with bulk methylation proportion *p*, in any given individual, the proportion of MM cells would be approximately *p*^2^, the proportion of MU cells 2*p*(1 −*p*), and the proportion of UU cells (1 −*p*)^2^ (see Methods for further details). For any given CpG, the likelihood of a particular methylation state being present in the original stem cell leading to clonal outgrowth would then be proportional to these probabilities.

To test this theory, we used a dataset consisting of 68 patients with acute myeloid leukemia (AML)^16^. We assumed that people with AML would experience high clonality, i.e., they should display methylation peaks close to the ratios of the true underlying allele distribution. As such, we calculated the observed ratios of peak heights in the AML dataset, along with expected peak heights under the HW assumption (see Methods for details). These results are presented in **Extended Data Fig. 1**, demonstrating that the HW assumption broadly aligns with the observed data.

Under the HW assumption, a theoretical clonal signal strength can be derived for any group of CpGs with a certain population mean, *p*, based on the extent to which the distribution of methylation values is expected to shift when a clone expands (see Methods for details). The result of this analysis is shown in the 1-dimensional heatmap in **Fig. 1b**, with the theoretical signal strength plotted across values of *p*. We expect the strongest signal for CpGs with a mean of approximately 0.33 and 0.66, with the signal decreasing to 0 at both extremes and a dip at 0.5.

### Datasets

For our subsequent analyses, we used two datasets. First, we used two longitudinal studies of cognitive aging: the Lothian Birth Cohort 1921 and the Lothian Birth Cohort 1936, hereafter referred to as the LBC dataset (see Methods for details). Briefly, participants were recruited in their eighth and seventh decades of life, with both cohorts undergoing comprehensive molecular profiling, including DNA methylation analysis, whole-genome sequencing (WGS), and targeted error-corrected sequencing. Unless specified otherwise, the ground-truth VAFs used for this study were taken from the targeted sequencing data. After quality control, the final dataset consisted of 71 participants with 108 CH measurements. Second, we used Generation Scotland (GenScot): a large (N=15,900) family-based cohort with paired methylation data and longitudinal health outcomes, consisting of adults aged 18-98.

### Simulated clonal expansion aligns with real data

To validate the theoretical framework of COMET, we created a forward simulation to predict the methylation distributions given i) a baseline representing a state of no clonal hematopoiesis, and ii) the sizes (i.e. VAFs) of the clones. To assess the concordance of this simulation and the actual data **(Fig. 1c)**, we selected a set of CpGs satisfying the assumptions outlined in **Fig. 1a** (see Methods for details) and with a population mean of ∼0.3, as per the theoretical maximum signal intensity shown in **Fig. 1b**. Next, we grouped participants from the LBC dataset based on VAF detected via targeted sequencing. We then predicted the shape of the methylation distributions for each VAF group using the methylation values from participants with the smallest clones (VAF 0-2%) as our baseline. As clonal expansions grow larger (increasing VAF), the methylation landscape progressively diverges from the normal state. The concordance between theoretical predictions (red) and observed data (blue) across all clone sizes validates that COMET accurately captures how clonal hematopoiesis reshapes the methylation landscape in blood.

Lastly, we investigated whether similar dynamics are also observed in mice. **Fig. 1d** shows the distribution of an equivalent set of CpGs from the mammalian methylation array (see Methods for details) in a young mouse (left panel) showing no evidence of clonal expansion, and an old mouse (right panel) showing extreme clonal dynamics. Together, **Fig. 1d** demonstrates that the collective behaviour of CpGs outlined above applies across species and platforms.

### Quantifying clonality from bulk methylation dynamics

Using these principles, we then developed a measure of the VAF in an individual based on bulk methylation data (COMET). First, CpGs were grouped based on their average methylation proportion (see Methods for details) so that HWE proportions could be calculated. On average, across the group of CpGs, any given clone would be expected to possess each underlying allele state in these HWE proportions. As demonstrated by **Fig. 1c**, the premise was that by observing many CpGs, the extent to which the methylation proportions diverged from the mean and towards a final state dictated by HWE proportions would indicate the level of clonality within a person.

Next, we developed a statistical method to quantify the level of CH for each person based on these CpG distributions. First, the expected number of CpGs in each allele state in the clone was calculated based on the HWE equations. Next, we assigned each CpG to one of these states, in these proportions, based on its closest value. This assigned state represents the best guess at the underlying allele state of the clone in that CpG. **Extended Data Fig. 2** illustrates an example with *p* = ∼0.3.

Lastly, the level of CH in a person can be quantified based on how far these CpGs have shifted from their original value. Briefly, we assumed that the difference from the original value (i.e. the mean in healthy people) represents the change in allele proportion due to the clone expanding. The COMET algorithm measures this change and returns a clonality score (referred to hereafter as COMET score) based on both the allele status of the clone and the allele status of the population of cells it is replacing (see Methods and **Extended Data Fig. 2** for details).

### Feature selection refines COMET performance

Next, as COMET relies on grouping CpGs with similar means, we aimed to evaluate its performance across a range of methylation values. We also hypothesized that not all CpGs would show evidence of clonal expansion, even within the same mean methylation range. For example, some CpGs might be biologically regulated and unable to significantly change their value. To estimate which CpGs showed the most evidence of clonal dynamics, we applied our CH estimation methodology to single CpGs. The principles behind the single-CpG application of the COMET algorithm are identical to those behind the single-person application described above. The only practical difference is that the distribution used to estimate clonality consists of values from a single CpG across many people, rather than values from many CpGs within a single person (see Methods for further details). To avoid biasing our estimates in the LBC cohort, we used GenScot to infer CpG clonality estimates. For each CpG, we compared the clonality estimates of the youngest people in the GenScot (those 20 years and younger) with those of the oldest (those 75 years and over), reasoning that older people tend to have a higher CH burden^3,7,8^. The CpGs with the highest difference (referred to hereafter as the CpG clonality score) are the ones that showed the most evidence of clonal dynamics developing over the lifespan.

**Fig. 2a** shows the results of applying this methodology to the LBC dataset across varying mean methylation values and CpG clonality scores. Each tile is coloured according to the R^2^ value that results from a simple linear regression of VAF (measured by targeted sequencing) against COMET scores for each sample. The best predictions result from CpGs with a mean methylation values of approximately 33% or 66% - as expected from the theoretical signal strength calculations - and a clonality score of greater than approximately 0.03-0.06. (Note that CpGs with poor clonality scores do not lead to accurate predictions - see **Extended Data Fig. 3** for comparison.)

**Figure 2.**
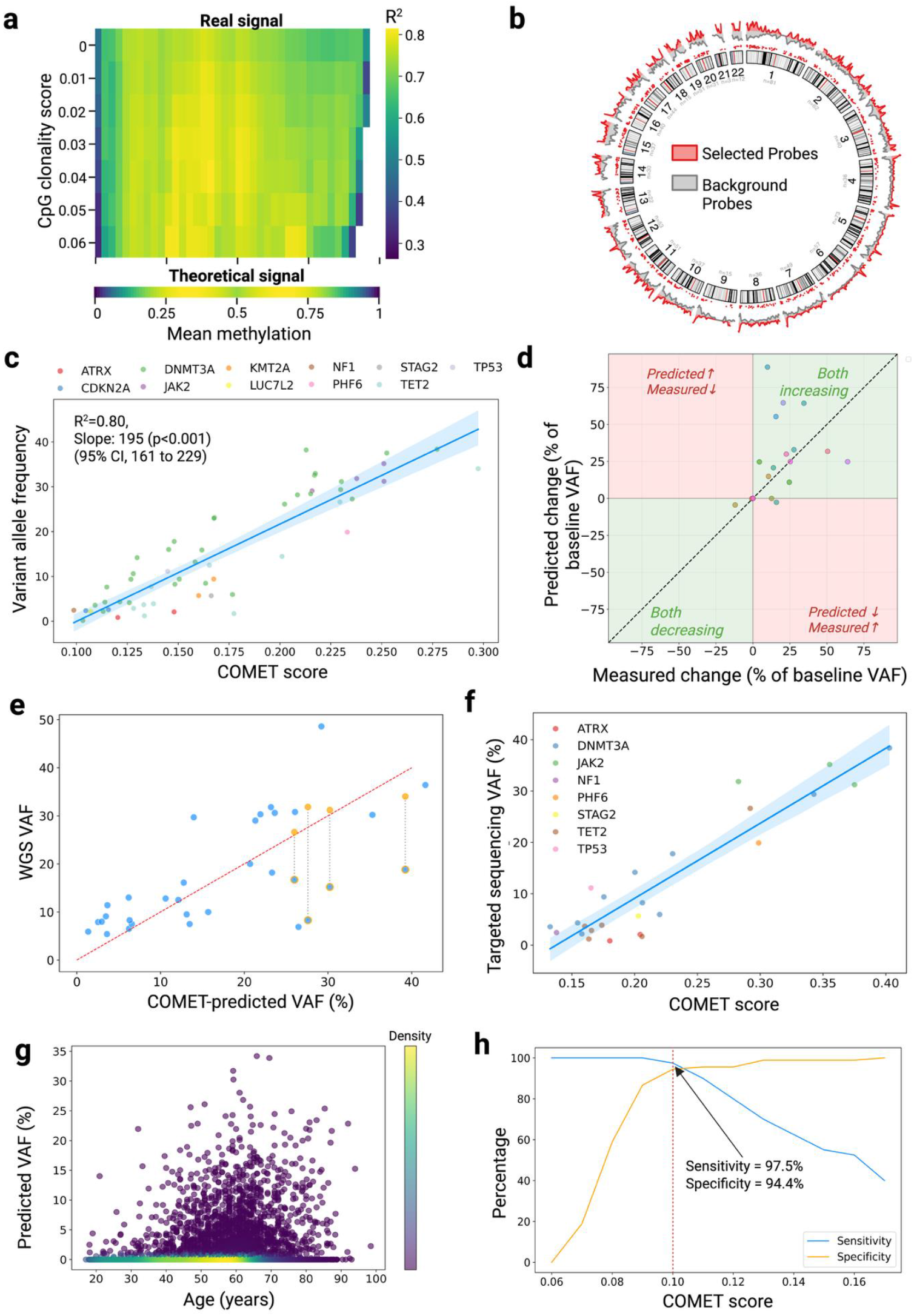
COMET Tracks Longitudinal VAF Changes, Detects Artifacts, and Differentiates Cases from Controls. **a**, Heatmap of strength of predictions using CpGs of various mean values (x-axis, binned to the value shown plus 0.05) and CpG clonality scores (y-axis, representing all CpGs with scores greater than the value shown). Strength of predictions measured by R^2^ value between COMET score and variant allele frequency (VAF), as measured by targeted sequencing. Missing segments reflect sample sizes <50. The theoretical signal strength is shown below (see Methods). **b**, Chromosomal location of selected CpGs for COMET (red) versus background probe distribution from Illumina Infinium HumanMethylationEPIC BeadChip (gray). **c**, Variant allele frequency (VAF), as measured by targeted sequencing, versus COMET score using the set of selected CpGs outlined in b). Samples consist of those with VAFs >0% and ≤50%, coloured according mutation detected. **d**, VAF as measured by targeted sequencing versus COMET predictions in samples with longitudinal measurements (i.e. across “waves” of the studies). VAF change shown relative to the targeted sequencing VAF at the first time point. **e**, VAFs, as measured by whole-genome sequencing, versus COMET predictions. Outlying points with an orange outline have corresponding measurements from the targeted sequencing (orange points) that align more closely to COMET predictions. The red dotted line shows y=x (i.e., perfect alignment). **f**, VAF, as measured by targeted sequencing, against COMET scores derived from a simplification of the COMET algorithm for CpGs on the X chromosome only. Only male samples are included in this analysis. **g**, Predicted VAFs in a large healthy cohort (N=15,900) aged 18-98. Negative predicted VAFs have been set to 0. **h**, Sensitivity and specificity of COMET using various COMET score thresholds to classify a case. Ground-truth cases were defined as any sample from the Lothian Birth Cohort with a detected VAF (i.e., >0%) from targeted sequencing. Young participants (aged 18-30) from Generation Scotland were used as controls.

We selected one of the best-performing combinations of mean methylation range and CpG clonality score that still retained a large number of CpGs (see **Supplementary Table 1** for full list and Methods for details). **Fig. 2b** shows the distribution of these CpGs across the genome. As can be seen, the CpGs are evenly distributed across chromosomes and follow the relative distribution of probes on the array, illustrating that the COMET criteria define a broad array of sites rather than a highly specific set of CpGs (as would be the case with machine-learning methods to predict CH from methylation).

**Fig. 2c** shows the results from these selected CpGs in samples from the LBC dataset with a detected CH mutation of ≤50% (see Methods for further details). VAF, as measured by targeted sequencing, is plotted against the COMET score for each sample. COMET score is highly correlated with VAF, with 80% of the total variance observed in VAF explained by our methylation-based predictor (R^2^ = 0.80). It is important to note that age explains essentially none of the variance observed in VAF values (R^2^ = 0.02, see **Extended Data Fig. 4**), so COMET does not simply reflect age. Lastly, points are coloured according to the specific CH mutation, showing that COMET performs similarly across all mutation types.

While COMET scores should theoretically broadly align with clonal proportions, **Fig. 2c** shows that, in practice, they require a slight adjustment. To convert COMET scores to predicted VAFs, we regressed the targeted sequencing VAFs on the COMET scores, using a mixed-effects model (random intercepts only) to account for the longitudinal measurements of some samples (slope = 195, 95% CI 161 to 229, p<0.001, intercept = -18.7). All subsequent predicted VAFs use this relationship to convert to VAF from the raw COMET score.

### Resolution improves by looking at longitudinal changes in VAF

As shown in **Fig. 2c**, COMET is best at detecting large absolute VAFs if performing a single measurement. However, resolution can be dramatically improved by measuring longitudinal changes within a subject. **Fig. 2d** shows the algorithm applied to each sample with longitudinal measurements, with the change shown relative to the targeted sequencing VAF at the first time point (one point omitted; see Methods for further details). COMET agrees with the directionality of change as measured by targeted sequencing in the vast majority of cases (2/2 negative changes and 14/16 positive changes). Across all longitudinal changes, the algorithm achieves a mean absolute error between first and last measurements of 1.7% - below the clinical VAF threshold of 2%, although noting that the longitudinal sample size is relatively small and that this result will change depending on the time between measurements.

### COMET can detect errors in sequencing-based methods and its outputs do not require manual curation

Next, we applied COMET to whole-genome sequencing (WGS) VAF measurements instead of targeted sequencing VAFs (**Fig. 2e**). As expected, the same trends exist, but with a lower correlation, given that WGS is lower-resolution than targeted sequencing. Additionally, **Fig. 2e** shows four large outliers (bottom-right of the plot, blue points with orange outline) for which COMET predicts a higher VAF than the WGS. For these points, there was a corresponding targeted sequencing measurement that confirmed the COMET predictions (solid orange points), showing the utility of the algorithm in detecting incorrect sequencing calls.

Similarly, **Extended Data Fig. 5** shows the results using targeted sequencing VAFs as in **Fig. 2c**, but this time with a) measured VAFs over 50% shown, b) samples with no detected CH shown, and c) outliers with a corresponding WGS measurement highlighted. Firstly, we can see that there is an instance of measured VAF exceeding 50%, which is corrected by COMET to a plausible VAF of ∼30%. In this instance, the variant in question was found to be the JAK2 V617F mutation, a locus that is widely reported for copy-neutral loss of heterozygosity (LOH) events in many myeloproliferative neoplasms (MPNs). (Indeed, many of these LOH events correspond to known diagnoses of MPN in the LBC dataset.) This is because COMET detects clonality across hundreds of CpG sites rather than examining specific mutations, and so is relatively unaffected by loss of heterozygosity in a certain gene. Secondly, we can see that three of the points with the highest discrepancy between COMET predictions and targeted sequencing have corresponding WGS measurements that more closely align with the COMET predictions.

Lastly, it is important to note that the sequencing-based VAFs in the LBC dataset represent the current gold standard for CH measurement. Not only are they from error-corrected targeted sequencing, but they are also manually curated to remove any dubious calls^12^. Prior to such manual curation, these sequencing-based methods return many calls for each sample, many of which are artifacts. (For a more comprehensive review of artifacts in CH calling, see Watson et al. (2020)^17^ and Robertson et al. (2022)^12^.) Without this expert manual curation, additional benefits of COMET become apparent. **Extended Data Fig. 6** shows the COMET score against the highest call for each sample from the non-curated targeted sequencing (i.e., the CH result that would be given for each person without dataset curation). Calls that were manually filtered out in the curated dataset are coloured in orange (i.e., artifacts), while correct calls are coloured in blue (i.e., real calls that also happen to be the highest call for a sample before dataset curation). Firstly, we can see that without data curation, 71% of the highest raw calls resulting from the targeted sequencing are likely artifacts. Secondly, unlike the real calls, we can see that the artifacts show minimal correlation with the COMET score. Again, this demonstrates the potential value the algorithm has to a) help filter sequencing-based CH results or b) to be used as a stand-alone method that does not require manual curation.

### Using only X chromosome CpGs improves COMET performance in males

COMET assumes that alleles at each CpG site are distributed according to Hardy-Weinberg Equilibrium. As males only have a single copy of the X chromosome, this scenario presents a special case in which this assumption is not needed; instead, the allele distribution is known definitively from the overall methylation proportion at each CpG. For example, a methylation proportion of 0.7 on an X chromosome CpG for a male means that 70% of their cells have their (only) allele methylated and 30% do not. By using these CpGs, the algorithm simplifies greatly (see Methods for details). Additionally, the clonality signal should be clearer as all methylation proportions move towards 0 or 1, whereas previously, the additional peak at 0.5 made small changes hard to detect.

**Fig. 2f** shows the results of applying this simplified version of COMET to X chromosome CpGs (see **Supplementary Table 2** for full list), using only the male samples from **Fig. 2c**. The correlation is very strong (R^2^ = 0.86) - slightly higher than using the method outlined in **Fig. 2c** - although the number of samples is quite low, so further data is needed to show that this is a robust improvement. **Extended Data Fig. 7** recreates the heatmap in **Fig. 2a** for the X chromosome CpGs in males, showing that the best correlations occur using CpGs with a mean of approximately 0.5.

### COMET strongly differentiates between cases and controls

As discussed above, some LBC samples without detected CH are likely false negatives, as evidenced by conflicting measurements between targeted and whole-genome sequencing. Additionally, given the known high prevalence of CH in older cohorts^3,7,8,^ such as the LBC, it is likely that at least some of the other outliers are also CH cases that were undetected by either sequencing method.

Because of this uncertainty, we decided to compare COMET predictions of CH cases with a set of young people aged 18-30 in the GenScot cohort. While there was no targeted sequencing data for this young cohort, our reasoning was that CH is very rare in younger adults^3,7,8^, and so they could more safely be used as controls than those in the LBC without detected CH.

First, we predicted CH in the entire GenScot cohort aged 18-98 **(Fig. 2g)**. While there is no ground truth for these samples, the predictions seemed sensible and aligned with expectations from prior literature. That is, CH is extremely rare in young people, but becomes relatively common in older adults. Also, it can again be seen that, while the predictions are age-related when comparing older and younger people, they are not so age-related that reliable predictions could be made solely on age, especially when looking within a set of people of similar ages (such as the LBC).

Next, we applied COMET to just those aged 18-30 and compared their scores with CH cases from the LBC (defined as those with any VAF detected via targeted sequencing, i.e., >0%). **Fig. 2h** shows the sensitivity and specificity of COMET in detecting a CH case using various score thresholds, with the optimum threshold achieving a sensitivity of 97.5% and specificity of 94.4%.

### Genome-wide association study of CH-associated genetic variants implicates TCL1A and NRIP1

Next, we performed a genome-wide association study (GWAS) to identify loci associated with methylation-predicted VAF in the larger GenScot cohort. **Fig. 3a** shows the resulting Manhattan plot annotated by mapped genes. The most significant hit was from the lead SNP rs2887399 on chromosome 22, which was mapped to the *TCL1A* gene - one of the most well-known CH-associated genes^19–22^. Two more hits were slightly below genome-wide significance (lead SNPs rs4890487 on chromosome 18 and rs2229742 on chromosome 21). rs2229742 mapped to gene NRIP1, which has also been recently implicated in CH^23,24^. (rs4890487 did not map to a gene.) Full lists of all implicated SNPs and lead SNPs can be found in **Supplementary Tables 3-4**.

**Figure 3.**
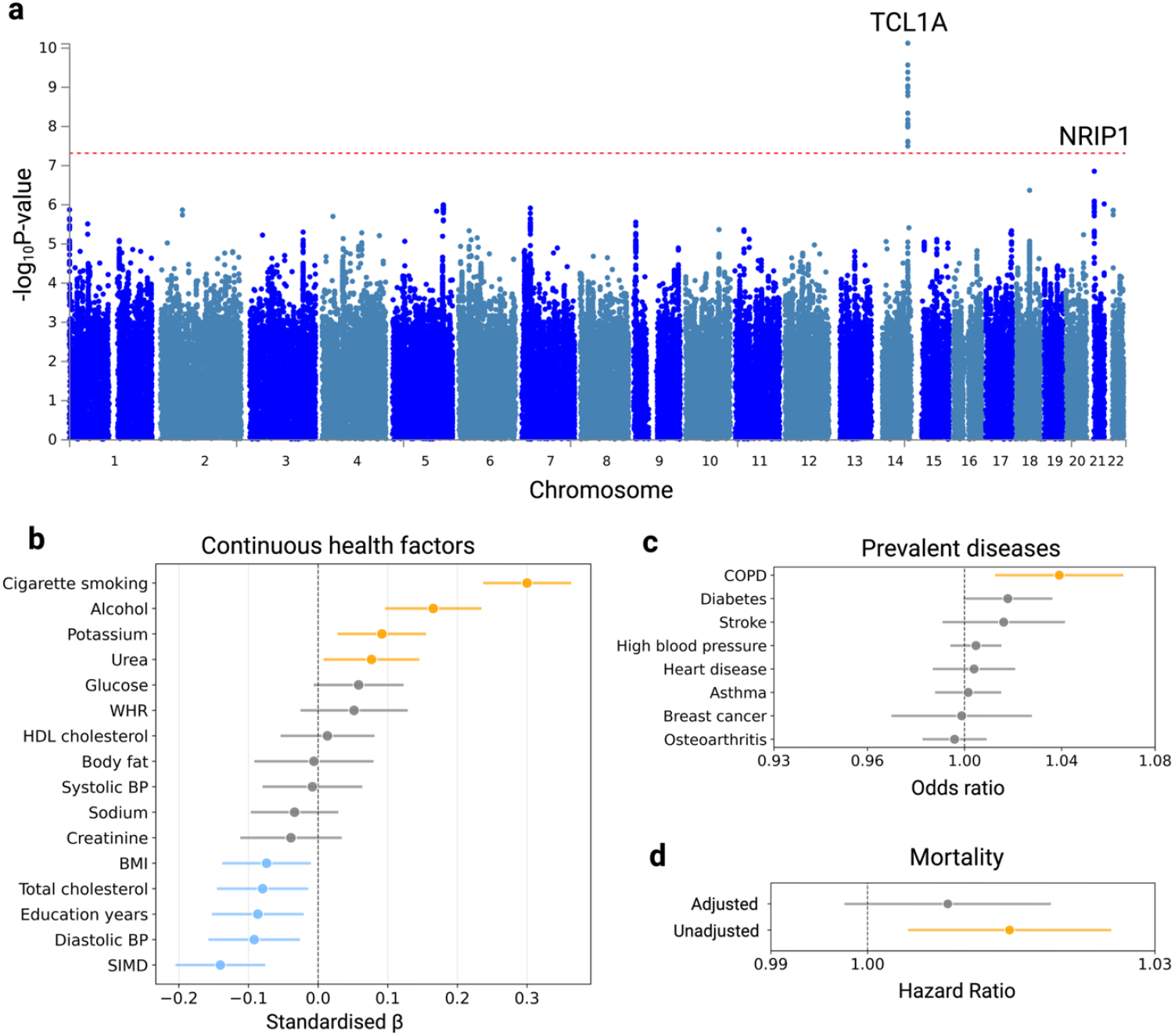
Biological validation of COMET. **a**, Manhattan plot from a genome-wide association study conducted on predicted CH values in the GenScot cohort. Red dotted line indicates genome-wide significance level. **b**, Associations of continuous lifestyle factors/health measures with predicted variant allele frequency (VAF, %). Dots denote standardised beta values (i.e. per 1 standard deviation increase in each variable) and horizontal lines represent 95% confidence intervals (CIs) estimated from linear regression models of the form. *VAF ∼ variable + sex + age*. Smoking is defined by a continuous variable that combines information on smoking status and quantity, and alcohol use is defined as the log of weekly units plus one (see Dabrowski et al.^18^ for further details). WHR: waist-hip ratio, BMI: body mass index, BP: blood pressure, SIMD: Scottish Index of Multiple Deprivation. Blue lines indicate FDR-significant (<0.05) protective associations, orange lines indicate significant adverse associations, and grey lines indicate non-significant associations. **c**, Associations of binary prevalent disease outcomes with predicted VAF. Dots denote odds ratios (per 1% increase in VAF) and horizontal lines represent 95% confidence intervals (CIs) estimated from logistic regression models of the form: *disease ∼ VAF + sex + age + smoking*. Smoking defined as in c). Blue lines indicate FDR-significant (<0.05) protective associations, orange lines indicate significant adverse associations, and grey lines indicate non-significant associations. **d**, Associations of predicted VAF with mortality hazard. Dot denotes hazard ratio (per 1% increase in predicted VAF) and horizontal lines represent 95% CIs estimated from a Cox proportional hazards model of the form: *hazard ∼ VAF+age+sex*. Smoking is also included as a covariate in the adjusted model. The orange line indicates a significant adverse association, and the grey line indicates a non-significant association.

### Predicted VAF is associated with lifestyle factors and health outcomes

Finally, we calculated associations between predicted VAF and a range of phenotypes in the GenScot cohort (**Fig. 3b-d**, see Methods for details and **Supplementary Tables 5-6** for full numerical results). In terms of lifestyle factors/health measures (**Fig. 3b**), there was a strong adverse association between predicted VAF and smoking (standardised beta value 0.30, 95% CI 0.23 to 0.36, FDR-adjusted p-value <0.001), which has also been seen in other studies^25,26^. We also see smaller adverse associations with alcohol use, urea and potassium levels, and protective associations with BMI, total cholesterol, education years, diastolic blood pressure, and Scottish Index of Multiple Deprivation (higher = less deprived). In terms of diseases (**Fig. 3c**), there was an association between predicted VAF and chronic obstructive pulmonary disease (odds ratio 1.04 per 1% increase in VAF, 95% CI 1.01 to 1.07), FDR-adjusted p-value=0.04), which also aligns with existing research^5^. Nominal associations were found with diabetes and stroke, although these did not quite pass FDR adjustment. Lastly, in terms of survival (**Fig. 3d**), there was an association between predicted VAF and all-cause mortality (hazard ratio 1.014 per 1% increase in VAF, 95% CI 1.004 to 1.025, p-value 0.006), although this weakened when adjusted for smoking (hazard ratio 1.008, 95% CI 0.998 to 1.019, p-value 0.12). However, the findings for disease and mortality outcomes should be interpreted with caution. Study power was quite low given the cohort was healthy, with low numbers of each outcome combined with low numbers of people with predicted CH.

## Discussion

We present COMET, a novel algorithm that quantifies CH burden using bulk DNA methylation data. This approach addresses key limitations of current CH detection methods by offering three major advantages: it requires minimal technical expertise, eliminates labor-intensive manual curation steps, and operates in a mutation-agnostic manner. This mutation-agnostic feature is particularly significant, enabling COMET to detect CH arising from driver mutations outside of standard gene panels and from large structural variants (e.g. loss of Y chromosome) that conventional approaches cannot capture. Additionally, COMET provides substantial cost savings compared to gene panel sequencing, with potential for even greater cost reduction through emerging platforms such as Illumina’s methylation screening arrays or targeted methylation sequencing.

COMET’s improvements position the method as an ideal screening tool, particularly for high-risk populations, such as those undergoing certain cancer therapies^10,27,28^. Positive cases identified through methylation screening could then undergo targeted sequencing for mutation characterization, creating a cost-effective two-tier diagnostic strategy.

COMET’s resolution dramatically increases when monitoring samples longitudinally. This improvement likely stems from the fact that there is variation in the methylation distributions of different people - even at the same VAF - but these differences are relatively consistent across time. Longitudinal sampling exploits this individual consistency by establishing person-specific baselines, thereby isolating true clonal changes from inter-individual methylation variability.

Several other studies have explored somatic epimutations as a promising clonal -tracking approach^14,29,30^, presenting an emerging area with increasing amounts of cohort-level single-cell data.

COMET’s utility theoretically extends beyond CH detection to any clinical scenario requiring clonal burden assessment. Applications include monitoring myeloproliferative neoplasms, acute myeloid leukemia, and multiple myeloma (using B-cell populations rather than whole blood). The approach could also potentially assess clonality in solid tumor biopsies, broadening its clinical relevance.

Key limitations include the inability to distinguish specific mutation types, though this trade-off enables detection of any CH clone regardless of driver mutations. While our method shows lower sensitivity than panel sequencing, it avoids false positives from sequencing artifacts, a major limitation of current approaches. The algorithm’s response to multiple large clones within individuals requires further investigation, though we anticipate reasonable robustness given that signals from different CpGs would partially cancel while others reinforce. Lastly, the sensitivity and specificity of COMET appear to be high, but ideally the algorithm would be tested with in an entirely separate dataset with age-matched cases and controls, both measured with the same type of methylation array and cross-validated via multiple methods.

Several refinements could enhance performance. Incorporating longitudinal data would improve accuracy by identifying trajectories inconsistent with birth-death clonal dynamics using our previously described LiFT filter^12^. Additionally, our algorithm assumes Hardy-Weinberg equilibrium at each CpG site, but incorporating sites that deviate from this distribution could enhance predictive accuracy if large reference datasets were available with either directly measured allelic methylation status or exceptionally high CH burden.

## Methods

### Datasets

To validate COMET, we used two main datasets. First, we used two longitudinal studies of cognitive aging: the Lothian Birth Cohort 1921 and the Lothian Birth Cohort 1936 (referred to as the LBC dataset). This cohort has been extensively described elsewhere^31^. Briefly, participants were initially assessed in childhood as part of the Scottish Mental Surveys conducted in 1932 and 1947, respectively, and subsequently recruited for follow-up studies beginning in their eighth and seventh decades of life. Both cohorts have undergone comprehensive molecular profiling, including DNA methylation analysis using Illumina methylation arrays (Illumina HumanMethylation450 BeadChip), whole-genome sequencing (WGS), and targeted error-corrected sequencing using a 75-gene panel (ArcherDX/Invitae). WGS data was processed as described in Roberston et. al (2019)^32^, and targeted sequencing was processed as described in Roberston et al. (2022)^12^. Unless specified otherwise, the ground-truth VAFs used for this study were taken from the targeted sequencing data. After quality control (see *Sample selection* below for details), the final dataset consisted of 71 participants with 108 CH measurements.

Second, we used Generation Scotland (GenScot): a large family-based cohort with paired methylation data and longitudinal health outcomes. This cohort has also been described extensively elsewhere^33^. Briefly, it consists of adults aged 18-98 recruited across Scotland from 2006 to 2011. Genome-wide DNA methylation was measured from blood samples using the Illumina Infinium HumanMethylationEPIC BeadChip. Samples were genotyped using the Illumina Human OmniExpressExome-8 v.1.0 and 8 v.1.2 BeadChips and processed using the Illumina Genome Studio software v.2011. Further details can be found in Dabrowski et. al (2024)^18^.

For **Extended Data Fig. 1**, we used AML data generated by Ferreira et al. (2016)^16^. DNA methylation was measured from blood samples using the Illumina HumanMethylation450 BeadChip. For **Fig. 1d**, we used mouse data obtained by the Mammalian Methylation Consortium^34^. DNA methylation was measured from blood samples using the mammalian methylation array. Further details can be found in Haghani et al.^34^.

### Methylation data processing

LBC DNA methylation data were processed using a normalization pipeline implemented in R using the minfi package^35^. Raw .idat files underwent quality filtering, removing samples with high mean detection p-values (>0.05), samples with >1% of probes having detection p-values >0.05, samples with low signal intensity, and samples with sex mismatches between predicted and recorded sex. Probes were filtered to exclude those with detection p-values >0.01 in >0.5% of samples, and cross-reactive probes were identified using the maxprobes package. The remaining data were processed using Noob background correction ^36^, followed by SWAN (Subset-quantile Within Array Normalization)^37^ to correct for probe-type bias and technical variation. Final beta and M-values were extracted for downstream analysis, with detection p-values >0.01 mean-imputed.

For the sensitivity and specificity analysis **(Fig. 2h)**, a subset of Generation Scotland was re-normalised in the same way as the LBC, described above, to ensure comparability between the two datasets. For all other analyses using Generation Scotland, the methylation data was processed as described in Dabrowski et al.^18^

The mouse methylation data was processed as described in Haghani et al.^34^. The AML dataset was processed as described in Ferreira et al. (2016)^16^.

### Using Hardy-Weinberg equilibrium to infer allele proportions

We hypothesised that the underlying proportions of allele combinations at a CpG with bulk methylation proportion *p*, in any given individual, should follow Hardy-Weinberg (HW) equilibrium:

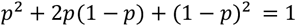

in which *p*^2^ is the proportion of MM cells, 2*p*(1 −*p*) is the proportion of MU cells, and (1 −*p*)^2^ is the proportion of UU cells. If CpG status does not affect whether a stem cell becomes a clone, then for any given CpG, the likelihood of a particular methylation state being present in the original stem cell will be proportional to these probabilities. **Extended Data Fig. 2** illustrates this idea. At 0% CH (no clonality), there is a methylation proportion of *p* = 0.3. In the example population of 10 cells, this means that roughly 10*p*^2^ = 10(0.3)^2^ = ∼1 cell is expected with state MM, 10(*p*)(1 −*p*) = 10(0.3)(1 −0.3) = ∼4 cells with state MU and 10(1 −*p*)^2^ = (1 −0.3)^2^ = ∼5 cells with state UU. This means that the probability of each state being present in the clone (and hence expanding) is ∼10% for MM, ∼40% for MU, and ∼50% for UU. These proportions will also determine the final peak heights at theoretical complete clonality.

### Validation of the Hardy-Weinberg assumption

To validate the HW assumption, we used an AML dataset. We assumed that people with AML would experience high clonality, i.e., they should display methylation peaks close to the ratios of the true underlying allele distribution. To calculate the observed peak heights in the AML dataset, we took the set of COMET CpGs and binned them into three categories for each sample: CpGs less than 0.33 were assigned UU, those between 0.33 and 0.66 were assigned MU, and those greater than 0.66 were assigned MM. To calculate the expected peak heights under HW, the HW proportions were calculated for each sample using sample-specific mean methylation values. These expected and observed proportions were then averaged across all samples included in the analysis.

### Calculation of optimal mean methylation values for maximising signal

#### Problem Setup

Given:

- Each CpG site has two alleles, each can be methylated (M) or unmethylated (U)
- Three possible genotypes: UU (0% methylation), MU (50% methylation), MM (100% methylation)
- Signal = maximum expected shift from starting value when a clone expands
- Assume underlying Hardy-Weinberg equilibrium (HWE)

#### Hardy-Weinberg Framework

Let *p* = population frequency of methylated alleles (M)

Under HWE, genotype frequencies are:

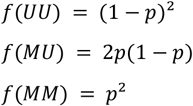

#### Signal Calculation

When a clone with variant allele frequency (VAF) = *v* expands, the observed methylation shifts, *β*_*obs*_, are the distance from the clone’s methylation value (0, 0.5, or 1) and the starting value, *p*:

- UU clone (methylation value = 0): *β*_*obs*_ = *v*(*p* −0) = *v*p
- MU clone (methylation value = 0.5): *β*_*obs*_ = *v*|0.5 −*p*|
- MM clone (methylation value = 1): *β*_*obs*_ = *v*(1 −*p*)

#### Expected Signal

The expected signal, E[S]:

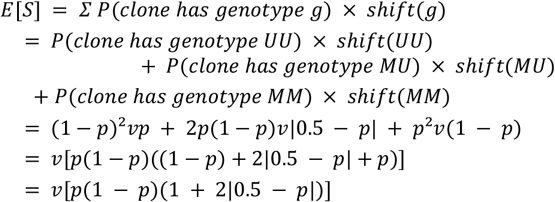

#### Optimization

To find the maximum for any given v, we need to maximize:

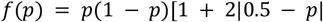

Breaking into cases:

**Case 1: *p*** ≤ **0. 5**

Here |0.5 −*p*| = 0.5 −*p*

f(*p*) = *p*(1 −*p*)[1 + 2(0.5 −*p*)]

= *p*(1 −*p*)[2 −2*p*]

= 2*p*(1 −*p*)^2^

= 2*p*(1 −2*p* + *p*^2^)

= 2*p* −4*p*^2^ + 2*p*^3^

Taking the derivative:

*f*′(*p*) = 2 −8p + 6p^2^

= 2(3*p*^2^ −4*p* + 1)

= 2(3*p* −1)(*p* −1)

Setting *f*′(*p*) = 0: *p* = 1/3 or *p* = 1

Since *p* ≤ 0.5, the critical point is *p* = 1/3

**Case 2: *p*** > **0. 5**

Here 0.5 −*p*| = *p* −0.5

(*p*) = *p*(1 −*p*)[1 + 2(*p* −0.5)]

= *p*(1 −*p*)[2*p*]

= 2*p*^2^(1 −*p*)

= 2*p*^2^ −2*p*^3^

Taking the derivative:

*f*′(*p*) = 4*p* −6p^2^

= 2*p*(2 −3*p*)

Setting *f*′(*p*) = 0: *p* = 0 or *p* = 2/3

Since *p* > 0.5, the critical point is *p* = 2/3

### Mathematical details of the clonal expansion model

To mathematically justify and validate the COMET algorithm, we developed a generative model to simulate the predicted distribution of bulk methylation data given an initial baseline distribution (*VAF* = 0%) and a series of clonal expansion events. This forward model allows us to test whether the theoretical framework underlying COMET can accurately reproduce observed methylation distributions across different clonal architectures.

The theoretical foundation assumes that as a clone with clonal fraction *c* = 2 · *VAF* expands, each CpG site’s methylation evolves according to:

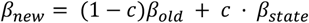

where *β*_*state*_ is sampled from one of three Hardy-Weinberg states: unmethylated-unmethylated (*β*_U*U*_ = 0), methylated-unmethylated (*β*_*M*U_ = 0.5), or methylated-methylated (*β*_*MM*_ = 1), with probabilities determined by the current methylation level *p*:

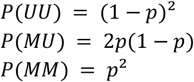

For a sequence of *n* clonal expansions, we can solve for the final methylation distribution analytically by enumerating the 3^*n*^ possible paths of Hardy-Weinberg outcomes. For each evaluation point *y* in the final distribution, we:

1. Consider each possible path (*s*_1_, *s*_2_, …, *s*_n_) where *s*_*i*_ ∈ {0.0, 0.5, 1}

2. Iterate backwards through the transformations to determine the original methylation value x_0_ that would produce y along this path

3. Compute the probability of this path occurring: *P*(*path*) = ∏*P*(*s*_*i*_|*x*_*i*™1_) using Hardy-Weinberg probabilities

4. Apply the Jacobian 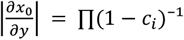 to account for the change of variables

5. Sum the contributions: *f*(*y*) = *Σ*_*paths*_ *f*_0_(*x*_0_) · *P*(*path*) · | *J* |

where *f*_0_ is the probability density of the original distribution. This analytical approach is mathematically exact but becomes computationally intractable for complex clonal architectures due to the exponential complexity of *O*(3^*n*^)

To facilitate rapid sampling of complex clonal variants and sequential expansion events, we implemented a Monte Carlo simulation that approximates the analytical solution. We sample *N* points from the baseline methylation distribution using kernel density estimation, then for each VAF in sequence:

1. Compute Hardy-Weinberg probabilities *P*(*UU*), *P*(*M*U), *P*(*MM*) for each sample using its current methylation value as *p*

2. Assign each sample to a methylation state via discrete sampling with these probabilities

3. Apply the mixture transformation:

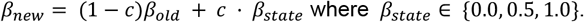

The final predicted methylation distribution is estimated via kernel density estimation on the transformed samples. With N=100,000 samples, this Monte Carlo approach converges to the analytical solution while remaining computationally efficient even for many sequential VAFs due to the linear complexity *O*(*n*)

While the simulated data broadly reflected the observed data, there were notable and systematic deviations between the base model predictions and empirical data. To account for these, we introduced two biologically motivated extensions:

#### Wright F-statistic

The base Hardy-Weinberg model assumes random assortment, but clonal populations in tissue exhibit non-random correlation. We incorporated Wright’s inbreeding coefficient *F* to model this clonal structure:

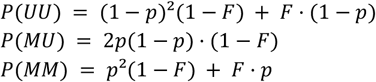

For visualization, we set *F* = 0.5 increases homozygous states (*UU* and *MM*) relative to heterozygous states (*M*U). Additionally, the discrete states {0.0, 0.5, 1.0} produce unrealistically sharp trimodal peaks in predicted distributions because DNA methylation maintenance with imperfect fidelity per cell division, meaning even monoclonal cells exhibit methylation heterogeneity due to stochastic maintenance errors. For visualization purposes, we replaced each discrete Hardy-Weinberg state with a continuous distribution in step 3 of the Monte Carlo algorithm:

- UU: *Beta*(*mode* = 0.1, κ = 15)
- MU: *Beta*(*mode* = 0.5, κ = 50)
- MM: *Beta*(*mode* = 0.9, κ = 7.5)

### Mathematical details of COMET

To illustrate the construction of the COMET algorithm, suppose we take the values of a group of CpGs in an individual with a mean *p* in the population. Suppose that one of those CpGs in this person has a methylation proportion of *x* and has been assigned MM (i.e., is one of the CpGs closest to 1). It has moved *x* −*p* from the mean, i.e., the proportion of methylated alleles has changed by *x* −*p* due to the clone expanding. Now, note that if the MM clone replaces a UU cell, then two extra methylated alleles will be gained in the population, while if it replaces a MU cell, then only one will be gained. (We can essentially ignore it replacing MM cells because it results in no change in the proportion of methylated alleles.) As such, if we let *c* = *number of clones, P*_*MM*_ = *proportion of >MM cells, P*_*M*U_ = *proportion of >MU cells, P*_*UU*_ = *proportion of MU cells*, and *x* −*p* = *the change in mean due to expansion of an MM clone*, then:

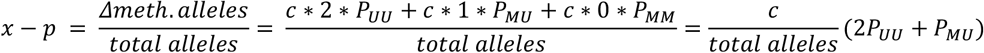

Note that 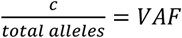, which we subscript with MM to denote it’s estimated from an MM clone,

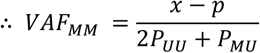

Equivalent expressions can be derived using this same logic for clones assigned UU:

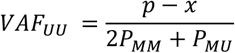

Similar expressions can also be formulated for MU clones, but for COMET, we ignore them as the signal from MU clones is the weakest, given that the maximum change in mean is the smallest. Similarly, in order to only use the strongest signal for each set of sites, we use the mean COMET score resulting from the UU estimates if the mean methylation value is above 0.5, and the score resulting from the MM estimates if the mean methylation value is below 0.5. Additionally, we first trim the highest 10% of COMET scores before calculating the mean to remove the influence of any highly outlying methylation values.

### Special case of COMET: X-chromosome CpGs in males

As males only have one copy of the X-chromosome, the COMET algorithm simplifies greatly when restricting to only X-chromosome CpGs in these participants. Specifically, the assumption of Hardy-Weinberg equilibrium is no longer necessary, and instead, the allele proportions are known definitively:

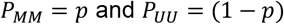

In such a case, the COMET equations simplify to:

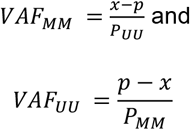

Note that we have lost the *P*_*MU*_ terms as this goes to zero, and we have also lost the factor of 2 from the *P*_*UU*_ /*P*_*MM*_ terms, as now we only gain one allele with each clone.

### Special case of COMET: calculating CpG clonality scores

The above algorithms have presumed that we are calculating scores for an individual by using the values from a set of CpGs with a similar mean. However, COMET is also applicable to calculating the clonality of individual CpGs by instead using the values of the CpG across many samples. The equations remain identical; it is simply the context that has changed. To summarise:

- Using COMET to calculate clonality in an individual: methylation values input are from many CpGs with a similar mean, taken from the same individual
- Using COMET to calculate clonality in a CpG: methylation values input are from the same CpG across many individuals

### Sample selection

Before the main analyses, we conducted a PCA on the methylation values in the LBC dataset. Samples clustered by sex **(Extended Data Fig. 8b)**. However, there was also a clear difference between “Sets” (i.e., experimental batches), with Sets 2 and 3 clustered tightly together, while samples from Set 1 were largely separate **(Extended Data Fig. 8b)**. This difference remained even after re-normalisation attempts, and so Set 1 was removed from all subsequent analyses. The final dataset consisted of 71 participants with 108 measurements.

For the longitudinal analysis shown in **Fig. 2d**, one sample was removed with a baseline VAF of exactly 0. Even with a small epsilon added to each denominator, the resulting relative change was too large for this sample to be clearly plotted with the other samples.

For the analysis of the AML dataset shown in **Extended Data Fig. 1**, nine samples were removed that had unimodal methylation distributions - i.e., did not satisfy the assumption that they approximated a completely clonal methylation distribution.

### CpG selection

#### Human data

As per the heatmap in **Fig. 2a**, CpGs were selected for COMET based on a combination of their mean methylation value and their clonality score. The final CpGs selected had a mean at age 0 of between 0.25 and 0.30 (i.e. the intercept resulting from a simple linear regression of methylation proportion versus age in the GenScot cohort), and a clonality score of greater than 0.04. Note that the mean values in the LBC are shown in the heatmap. CpGs with an absolute mean change (i.e. slope) of >0.05% year were excluded as Hardy-Weinberg equilibrium proportions could not be reliably calculated. CpG clonality score was calculated by applying the CpG-specific COMET algorithm to younger (aged <=20) and older (aged >=75) GenScot participants separately and taking the difference (older minus younger). The reasoning behind this was that CH is known to be highly prevalent in older people but relatively rare in younger people, and so CpGs that showed a difference in clonality between these two subsets would likely be implicated in CH.

#### Mouse data

For the mouse data in **Fig. 1d**, the same CpGs could not be used as in the human analyses as a) the array used for the mouse data was the Mammalian Methylation Array and so shared relatively few CpGs with the human 450k array and b) the CpGs satisfying the assumptions outlined in **Fig. 1a** are not necessarily the same between human and mouse. As such, different CpGs had to be selected, but the methods outlined for human data above translated poorly to the smaller sample size of the mouse data. Instead, CpGs were selected based on those that showed the highest variance change with age (comparing the variance of mice over and under the median age), and with an intercept of between 0.2 and 0.3 (resulting from a simple linear regression of methylation proportion versus age).

### GWAS

The GWAS was performed using the same dataset and similar methods as outlined in Dabriowski et. al^18^. Briefly, genotype–phenotype association analyses were performed using a linear mixed model GWAS implemented in fastGWA GCTA^38^. A sparse GRM approach was used to adjust for sample-relatedness. Overall, 15,871 overlapping individuals with non-missing genotype and phenotype data were included. Variants with MAF < 0.01 or a missingness rate > 0.1 were excluded from the analysis. Genomic risk loci were defined around significant variants (P<5×10^−8^) and included all variants correlated (R^2^>0.1) with the lead variant. All coordinates in this study were based on human reference genome assembly GRCh37/hg19 (http://www.ncbi.nlm.nih.gov/assembly/2758/). Functional analysis of the GWAS results was performed using FUMA^39^ with default parameters and the UK BioBank release 2b 10k White British reference population.

## Data availability

### Lothian Birth Cohorts data

Data from the Lothian Birth Cohorts is available upon request. Further information can be found here: https://www.ed.ac.uk/lothian-birth-cohorts.

### Generation Scotland Wave 3 data

According to the terms of consent for Generation Scotland participants, access to data must be reviewed by the Generation Scotland Access Committee. Applications should be made to.

### Mouse data

The mouse dataset used was created by the Mammalian Methylation Consortium^34^ and is publicly available on the Gene Expression Omnibus under accession number GSE223748.

### Acute myeloid leukemia (AML) data

The AML data used was described in Ferreira et al. (2016)^16^ and is publicly available on the Gene Expression Omnibus under accession number GSE62298.

## Ethics

### Lothian Birth Cohorts

Ethics permission for the LBC1921 was obtained from the Lothian Research Ethics Committee (Wave 1: LREC/1998/4/183; Wave 2: LREC/2003/7/23; Wave 3: LREC1702/98/4/183) and the Scotland A Research Ethics Committee (Wave 4: 10/S1103/6; Wave 5: 10/MRE00/87). LBC1936 ethical approval was obtained from the Multicentre Research Ethics Committee for Scotland (Wave 1, MREC/01/0/56), the Lothian Research Ethics Committee (Wave 1, LREC/2003/2/29), and the Scotland A Research Ethics Committee (Waves 2-5, 07/MRE00/58). All participants provided written informed consent.

### Generation Scotland

All components of GS received ethical approval from the NHS Tayside Committee on Medical Research Ethics (REC reference no. 05/S1401/89). GS has also been granted Research Tissue Bank status by the East of Scotland Research Ethics Service (REC reference no. 20-ES-0021), providing generic ethical approval for a wide range of uses within medical research.

## Code availability

All code used for the analysis (conducted in Python 3.11.6) is available at https://gitfront.io/r/scrofts/TXcXutEt29G9/COMET/.

## Extended Data

**Extended Data Figure 1.**
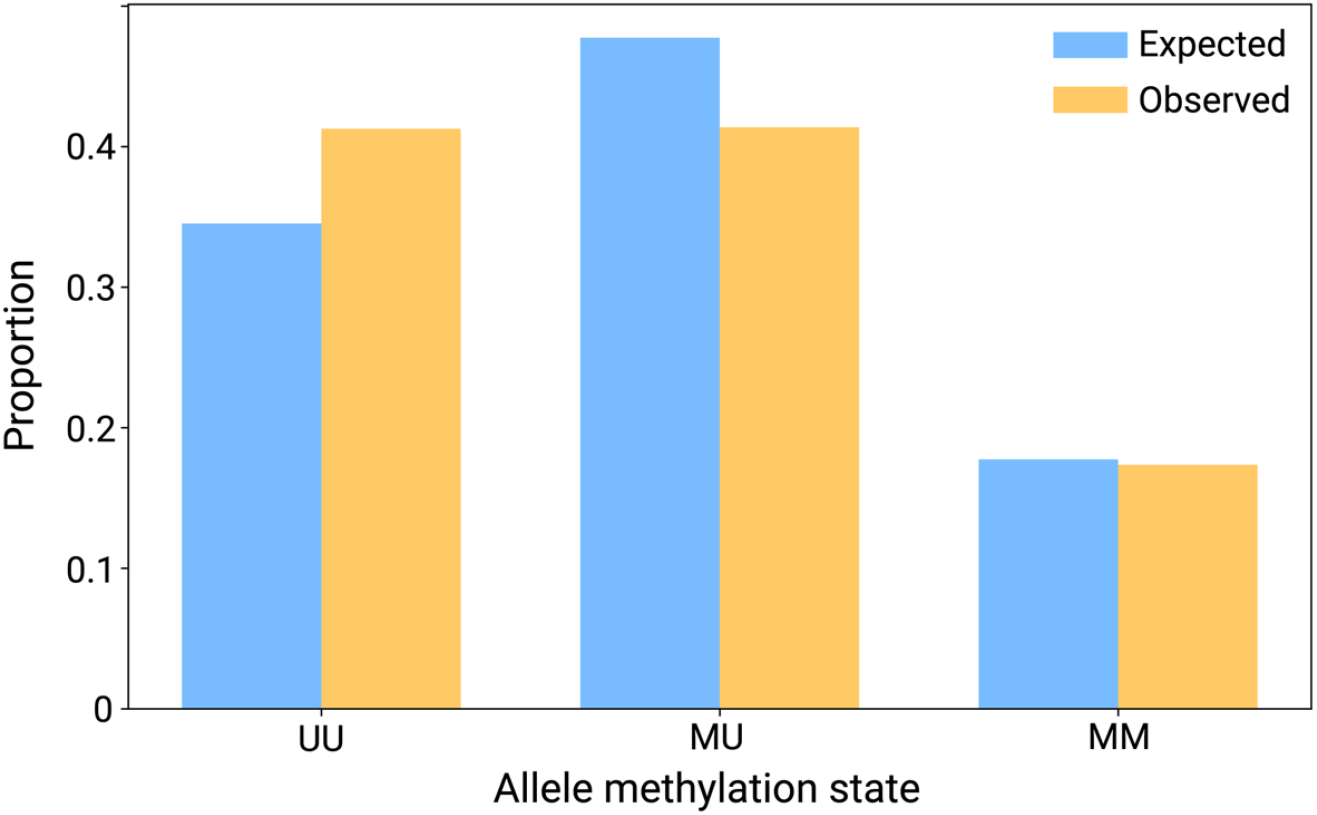
Observed versus actual proportions in each methylation state in patients with acute myeloid leukemia (AML) Observed versus expected (under the Hardy-Weinberg assumption) allele proportions in patients with AML. As these patients are presumed to exhibit very high clonality, the observed proportions are used to gauge the “final state” of clonality, i.e. the true underlying allele proportions.

**Extended Data Figure 2.**
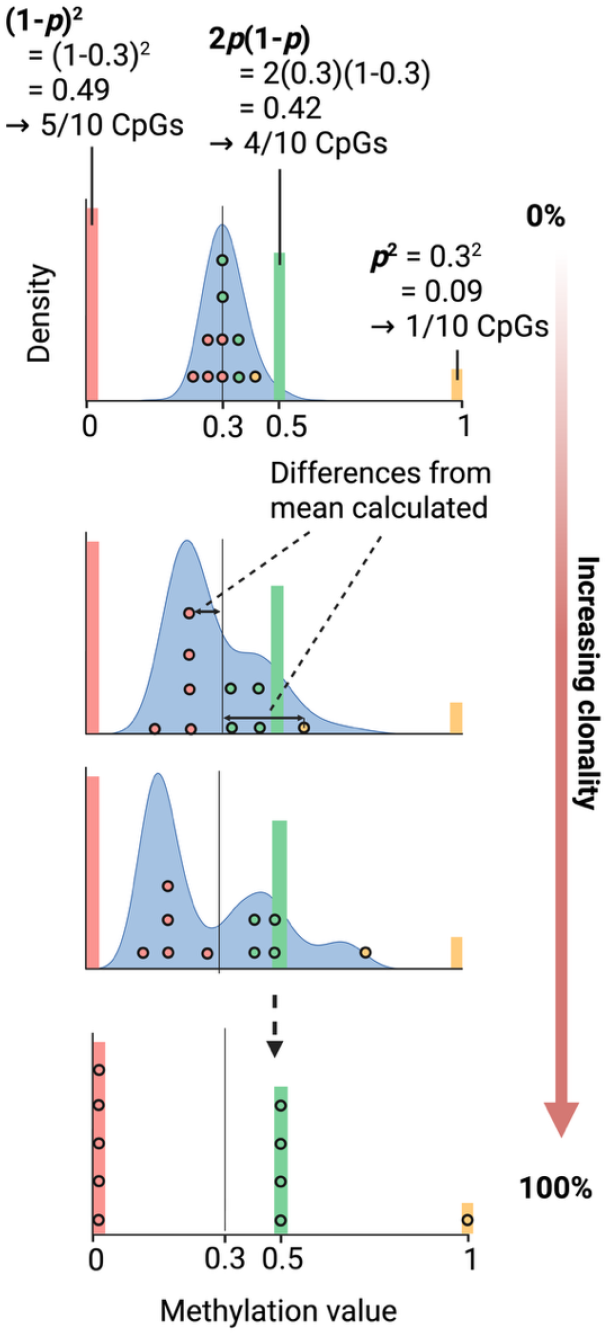
Diagrammatic overview of the COMET algorithm. Overview of the COMET algorithm, using the example of sites with a population mean of ∼0.3. 1) A set of CpGs with similar mean values is selected, 2) Hardy-Weinberg Equilibrium (HWE) proportions are calculated based on this mean value, 3) CpGs are assigned to each allele state based on their closest value, and in the proportions given by HWE, 4) Distances from the mean value are calculated for each CpG, 5) Based on how far a CpG value has moved from the mean, taking into account its assigned allele state and the underlying allele distribution in the population, a score is calculated (see Methods for further details).

**Extended Data Figure 3.**
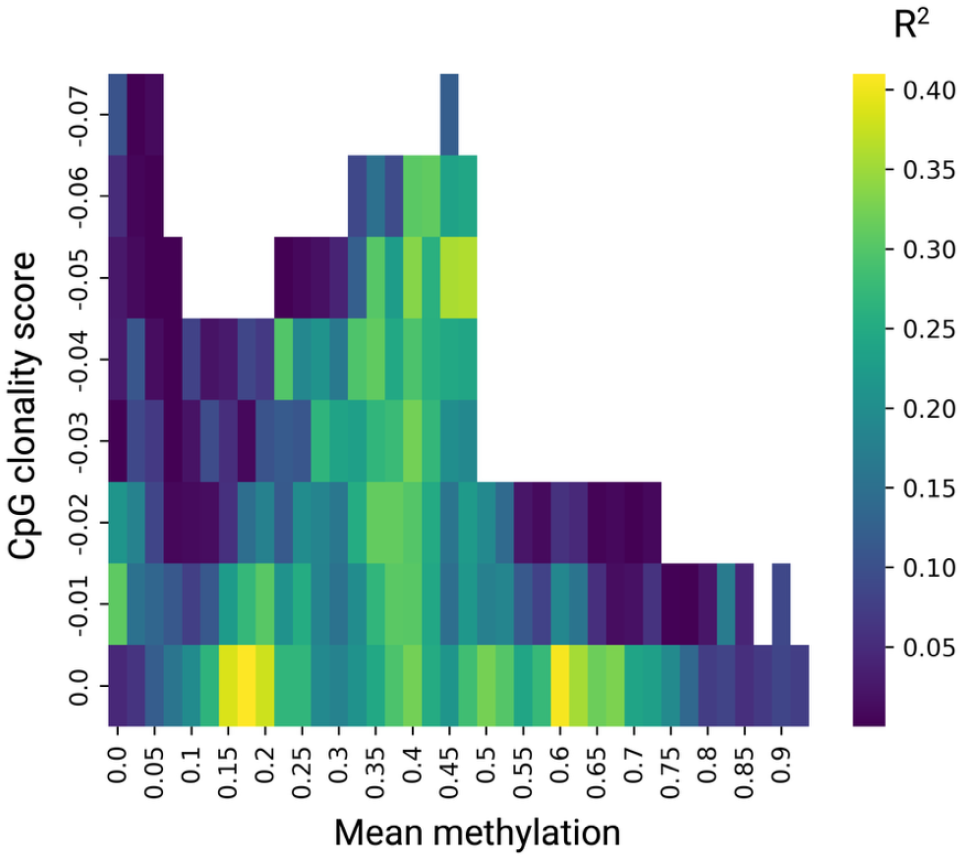
Heatmap of strength of predictions using CpGs with negative clonality scores. Heatmap of strength of predictions using CpGs of various mean values (x-axis, binned to the value shown plus 0.10) and clonality changes with age (y-axis, representing all CpGs with scores *less* than the value shown), using the COMET algorithm. Strength of predictions measured by R^2^ value between COMET score and variant allele frequency (VAF), as measured by targeted sequencing.

**Extended Data Figure 4.**
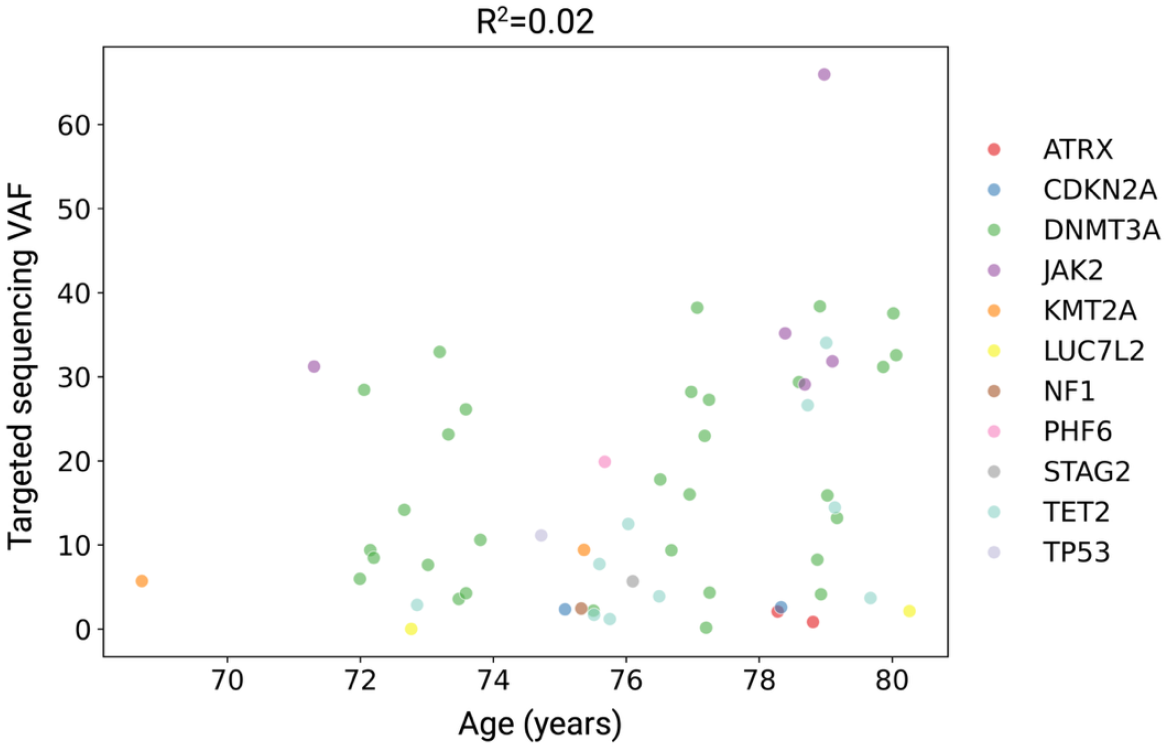
Association between VAF and age in the Lothian Birth Cohorts dataset. VAF (as measured by targeted sequencing) versus age (years). Points coloured by mutation detected.

**Extended Data Figure 5.**
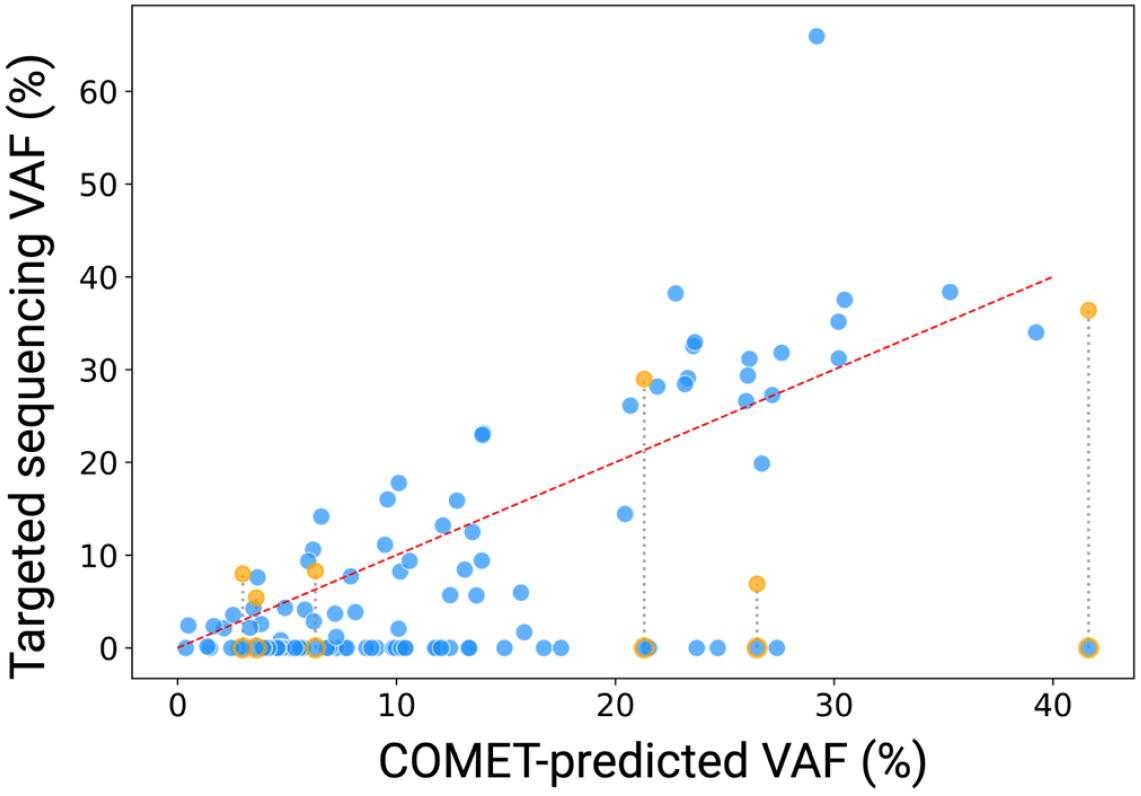
COMET can highlight potential errors with targeted sequencing. VAFs, as measured by targeted sequencing, versus COMET predictions. Samples include those for which no variant was detected (VAF=0%) and those for which a VAF of over 50% was measured. Outlying points with an orange outline have corresponding measurements from whole-genome sequencing (orange points) that align more closely with COMET predictions. The red dotted line shows y=x (i.e. perfect alignment).

**Extended Data Figure 6.**
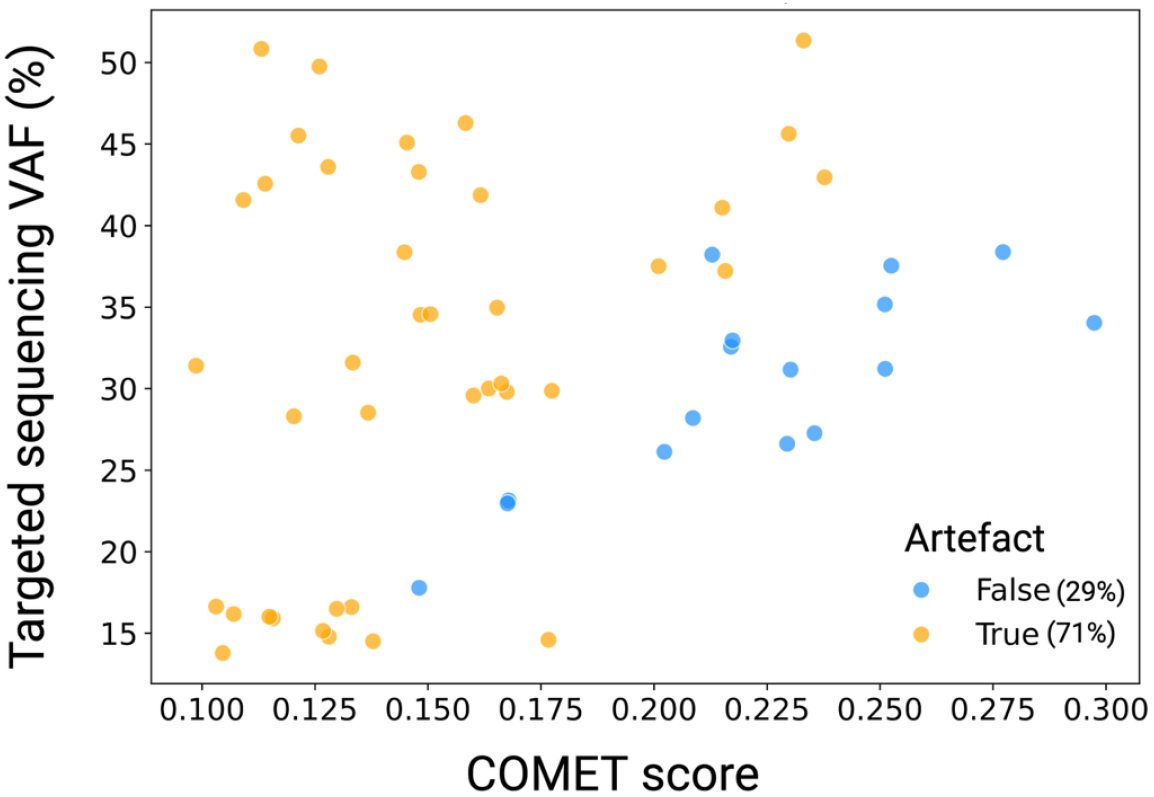
COMET correlates only with real sequencing calls. VAFs as measured by non-curated targeted sequencing (i.e. simply taking the highest call that is output for each sample) against raw COMET scores. Calls in samples for which the highest non-curated call happens to be the same as the curated call are classified as non-artifacts and coloured blue, while other calls are classified as artifacts and coloured orange.

**Extended Data Figure 7.**
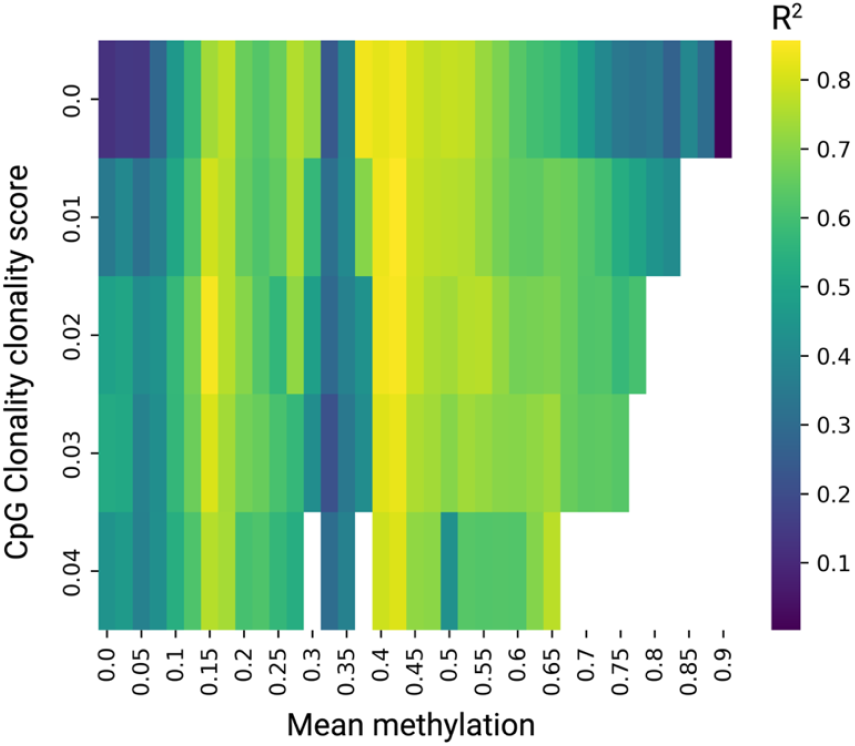
Heatmap of strength of predictions using X-chromosome CpGs in males. Heatmap of strength of predictions using CpGs of various mean values (x-axis, binned to the value shown plus 0.10) and clonality changes with age (y-axis, representing all CpGs with scores greater than the value shown), using the COMET algorithm adapted for use on X-chromosome CpGs in males only. Strength of predictions measured by R^2^ value between COMET score and variant allele frequency (VAF), as measured by targeted sequencing.

**Extended Data Figure 8.**
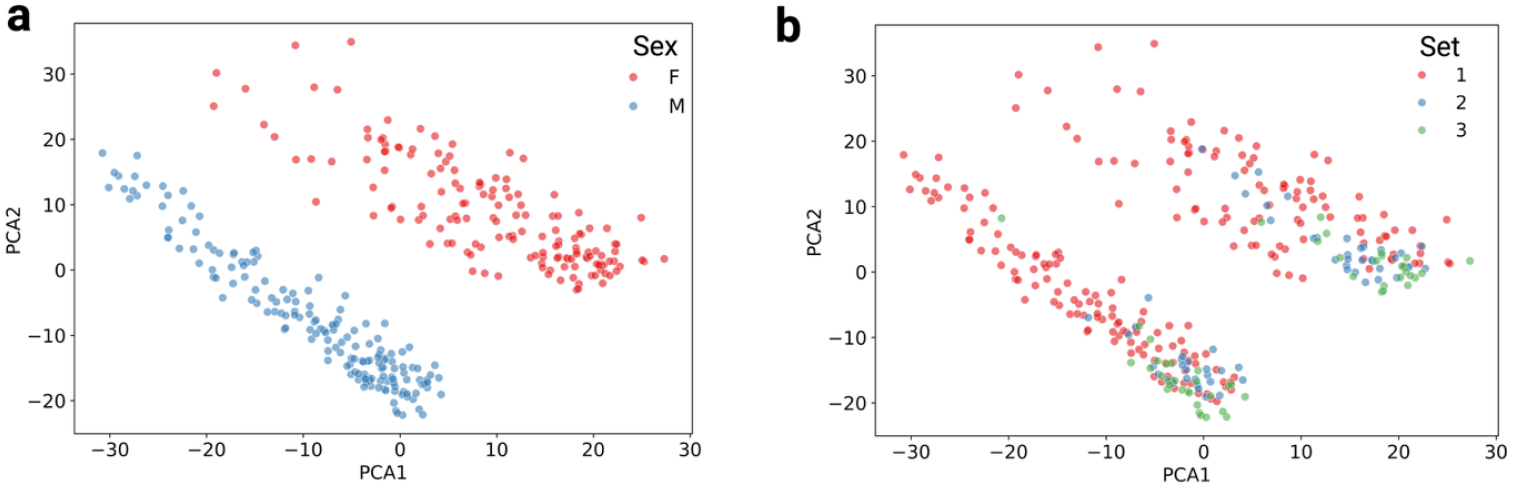
PCA of Lothian Birth Cohorts methylation data. **a**, PCA plot of all methylation values from the Lothian Birth Cohorts, coloured by sex. PCA 1 accounts for 13.9% of the total variance in methylation values, and PCA 2 accounts for 11.9%. **b**, PCA plot of all methylation values from the Lothian Birth Cohorts, coloured by “Set” (i.e. processing batch).

## Acknowledgements

We thank all Lothian Birth Cohort (LBC) and Generation Scotland (GS) study participants and research team members who have contributed, and continue to contribute, to ongoing studies. S.J.C.C. was supported by the Wellcome Trust Hosts, Pathogens and Global Health Programme (grant number grant.226831/Z/22/Z). T.C. and K.K. were funded by the Mayo Clinic Robert and Arlene Kogod Center on Aging. K.K. is funded by the Division of Hematology, Mayo Clinic, Rochester. The LBC1921 was supported by the UK’s Biotechnology and Biological Sciences Research Council (BBSRC), The Royal Society, and The Chief Scientist Office of the Scottish Government. LBC1936 is supported by the BBSRC, and the Economic and Social Research Council [BB/W008793/1], Age UK (Disconnected Mind project), the Milton Damerel Trust, and the University of Edinburgh. Methylation typing of LBC1921 and LBC1936 was supported by Centre for Cognitive Ageing and Cognitive Epidemiology (Pilot Fund award), Age UK, The Wellcome Trust Institutional Strategic Support Fund, The University of Edinburgh, and The University of Queensland. The whole genome sequencing of LBC1921 and LBC1936 was funded through an institutional award to the Roslin Institute from the BBSRC. GS received core support from the Chief Scientist Office of the Scottish Government Health Directorates (CZD/16/6) and the Scottish Funding Council (HR03006). DNA methylation profiling of the GS samples was carried out by the Genetics Core Laboratory at the Wellcome Trust Clinical Research Facility, Edinburgh, Scotland and was funded by the Medical Research Council UK, the Brain & Behavior Research Foundation (ref. 27404) and the Wellcome Trust (Wellcome Trust Strategic Award ‘STratifying Resilience and Depression Longitudinally’ ((STRADL) ref. 104036/Z/14/Z)).

